# Sex-dependent polygenic effects on the clinical progressions of Alzheimer’s disease

**DOI:** 10.1101/613893

**Authors:** Chun Chieh Fan, Sarah J. Banks, Wesley K. Thompson, Chi-Hua Chen, Linda K. McEvoy, Chin Hong Tan, Walter Kukull, David A. Bennett, Lindsay A. Farrer, Richard Mayeux, Gerard D. Schellenberg, Ole A. Andreassen, Rahul Desikan, Anders M. Dale

## Abstract

Sex differences in the manifestations of Alzheimer’s disease (AD) are under intense investigations ^1,2^. Despite the emerging importance of polygenic predictions for AD ^3–8^, the sex-dependent polygenic effects have not been demonstrated. Here, using a sex crossover analysis, we show that sex-dependent autosomal genetic effects on AD can be revealed by characterizing disease progress via the hazard function. We first performed sex-stratified genome-wide associations, and then applied derived sex-dependent weights to two independent cohorts. Sex-matched polygenic hazard scores (PHS) have significantly stronger associations with age-at-disease-onset, clinical progressions, amyloid depositions, neurofibrillary tangles, and composite neuropathological scores, than sex-mismatched PHS, independent of apolipoprotein E. Models without using hazard weights, i.e. polygenic risk scores (PRS), have lower predictive power than PHS and show no evidence for sex differences. Our results indicate revealing sex-dependent genetic architecture requires the consideration of temporal processes of AD. This has strong implications not only for the genetic underpinning of AD but also for how we estimate sex-dependent polygenic effects for clinical use.

---

Sex, as both an endogenous and an exogenous factor modulating human biology, has a ubiquitous impact on the pathogenesis of complex diseases ^9^. Evidence on sex-dependent clinicopathological progressions of Alzheimer’s disease (AD) is just beginning to emerge ^2^. Compared to men, women show later manifestation of verbal memory deficits ^10^, faster decline after disease onset ^11^, and some differences in neuropathological characteristics, such as tau tangle density ^1,12^. Results from studies on incidence rate and prevalence are less consistent ^13,14^, yet women are often reported to have increased incidence of AD in older ages ^15^ and higher prevalence ^16^. Given this unmet need for better understanding of sex differences in AD, we wanted to investigate the potential for a sex-dependent genetic architecture of AD. Despite evidence suggesting that AD is highly polygenic, with a heritability as high as 79 percent ^17^, so far only apolipoprotein E (*APOE* e4) has been found to have a differential impact on age-at-onset between men and women ^18,19^. Sex-dependent differences in polygenic effects remain unresolved.

This sex-agnostic status quo is particularly problematic for disease prediction based on polygenic effects. By aggregating the estimated regression weights of autosomal single-nucleotide-polymorphisms (SNPs) from genome-wide association studies (GWAS), polygenic scores have been used to assist in several important clinical functions, including disease prediction ^20^, risk stratification ^21^, enriching clinical trials ^6,22^, and facilitating disease screening ^23^. However, because the standard practice in GWAS treated sex as a confounding factor for autosomal effects, the basis of polygenic scores, the estimated odds ratios, are devoid of sex-dependent effects. Given the complexity of the moderating effects of sex on disease etiology ^9^, applying sex-agnostic polygenic scores may produce substantially biased risk quantifications. Such scores could underestimate the genetic risk of AD for women, since *APOE* e4, one of the most well-established risk factors for AD, has stronger effects on AD onset among women than among men ^19^. Given the heightened awareness of utilizing polygenic effects beyond *APOE* as biomarkers for AD ^5,22,24–27^, understanding the sex-dependent polygenic effects for AD is imperative for their application to clinical settings.

To investigate whether there are sex-dependent polygenic effects in addition to *APOE*, we performed a sex *crossover* study (Methods and Figure 1) – we derived polygenic scores from separate GWAS on men and women in the training cohorts (Alzheimer’s Disease Genetifc Consortium, ADGC, n = 17855; See Methods and Table 1), and then applied each of the sex-dependent regression weights to both men and women in independent cohorts (National Alzheimer’s Coordinate Center cohort, NACC, n = 6076; Religious Orders Study and Rush Memory and Aging Project, ROSMAP, n = 599) to determine if there were a differential performance in predicting AD. Importantly, the regression weights used as the basis for the scores were based on Cox regressions, thereby capturing differences in clinical progression between men and women as hazard functions.

**Table 1.** Characteristics of training samples and independent validating cohorts.

|  | Training samples |  | Independent testing cohorts |  |  |  |
| --- | --- | --- | --- | --- | --- | --- |
|  | *ADGC\** |  | *NACC* |  | *ROSMAP* |  |
|  | Men | Women | Men | Women | Men | Women |
| **Total N** | 7158 | 10697 | 2628 | 3448 | 220 | 379 |
| **Age - years (SD)** | 75.4 (7.7) | 75.9 (8.2) | 78.6 (9.4) | 79.1 (9.8) | 86.4 (6.3) | 89.4 (6.2) |
| **AD cases/events** | 42.7% | 47.6% | 52.3% | 41.5% | 37.7% | 43.8% |
| **APOE ε4 carriers** | 40.9% | 43.3% | 40.9% | 37.9% | 29.5% | 28.4% |

\* Excluded any overlapping samples with NIA ADCs and ROSMAP.
† NP – neuropathology. NP samples means number of samples with post-mortem neuropathology examinations.

**Figure 1.**
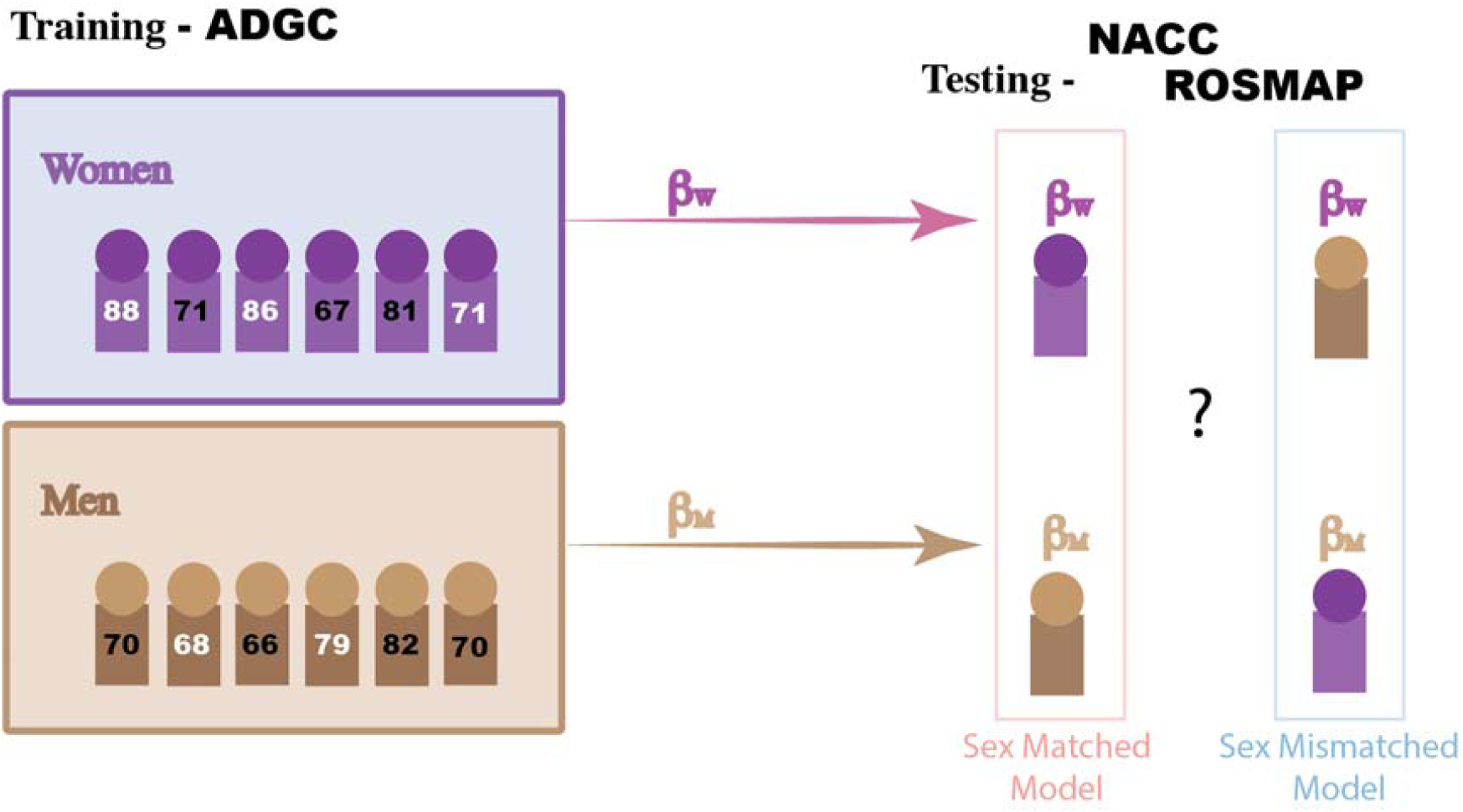
Flow chart of our sex crossover analysis.

## Methods

### Study Design

The crossover analysis is illustrated in Figure 1. First, we performed sex-stratified genome-wide analyses on age-at-onset of AD, using imputed genotypes and phenotypic data from the Alzheimer’s Disease Genetic Consortium (ADGC) ^28–30^. To ensure independence between the training and validation cohorts, we performed an extensive check on potential sample overlap and removed any overlapping individuals from the training data. The final training data included 7158 men and 10697 women (Table 1). Genome-wide Cox regression analyses were performed on men and women separately to obtain sex-dependent weights. Detailed descriptions of the analytical methods can be found in the following section and in the Supplemental Materials.

After obtaining the sex-dependent Cox regression weights for each autosomal SNP from the ADGC data, we applied these weights to two independent cohorts (Table 1), generating men-dependent polygenic hazard score (mPHS) and women-dependent polygenic hazard score (wPHS) for every participant. Thus, we can compare whether sex-matched models (mPHS on men and wPHS on women) has better predictive power than sex-mismatched models (wPHS on men and mPHS on women), as a cross-over comparison (Figure 1).

The first independent cohort was obtained from the National Alzheimer’s Coordinate Center (NACC). NACC recruits case series as a nationwide recruiting effort funded by National Institute of Aging, involving clinical centers across United States. Given the longitudinal design of NACC, we examined whether sex-matched PHS predicted dementia onset better than sex-mismatched PHS. The cohort characteristics of NACC can be found in Table 1.

The second independent cohort was the Religious Orders Study and Rush Memory and Aging Project (ROSMAP). ROS and MAP are two community-based cohort studies that enrolled individuals without dementia, all of whom agreed to longitudinal follow-up and organ donation, enabling us to examine the distribution of neuropathology among as a function of sex-specific PHS. All participants signed an informed consent, Anatomic Gift Act, and repository consent allowing their data to be shared. Both studies were approved by an Institutional Review Board of Rush University. Details of the studies, generation of genomic data, and neuropathologic data collection have been previously reported ^31,32^. We investigated whether sex-matched PHS has stronger associations with neuropathology in the brain than sex-mismatched PHS. For those who have both genotyping data and autopsy results were included in this analysis (n = 599). Detailed characteristics of ROSMAP can be found in Table 1.

For comparison purposes, we also examined the performance of polygenic risk scores (PRS) in the same manner as described above, except using weights from logistic regressions while controlling for age-at-ascertainment. This is intended to investigate the benefit of using Cox regressions in contrast to the standard GWAS approach.

### Estimating sex dependent hazards for autosomal SNPs

To obtain sex-dependent weights for each SNP, we fitted genome-wide Cox regression models on men and women separately. This stratified approach was intended to capture sex-specific effects from autosomal SNPs without explicitly modeling interaction terms. This stratified approach also allows for differences in the shape of the baseline hazard function between men and women. As noted in prior studies on sex-dependent genetic effects ^9^, although the total sample size for GWAS is thus reduced by half, stratified models are computationally simple and avoid the need for additional assumptions on the nature of sex interactions. Furthermore, hazard ratio estimation is facilitated by utilizing Martingale residuals under null ^33^:

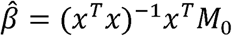

where x is the mean centered genotype dosage and M0 is the Martingale residuals of the null model. More detailed discussion about the hazard estimates from case-control study can be found in Supplemental Materials.

For ADGC data, we used the age-at-onset as the time-to-event and the age-at-last-visit as the censoring time for Cox regression while controlling for dosages of *APOE* e2 and e4, the first five genetic principal components, and indicators of recruiting sites. After filtering (minor allele frequencies greater than 1 percent, in Hardy-Weinberg equilibrium, missing rate less than 10 percent, located outside of *APOE* or major histocompatibility complex regions), 6,784,887 imputed SNPs were included in our analyses. The resulting men- and women-derived hazard ratios were used to generate the corresponding sex-dependent PHS. For comparison purposes, we also performed standard GWAS with logistic regressions for the same 6,784,887 SNPs. All covariates are the same in the models except age-at-ascertainment is now treated as one of the covariates. The estimated sex-dependent odds ratios were then used to generate the corresponding polygenic risk scores (PRS). Because our focus was on polygenic effects over and above the effects of *APOE*, we excluded any SNPs located within *APOE* region when we calculated all polygenic scores.

### Deriving polygenic hazard scores and polygenic risk scores

The polygenic scores are the product sum of GWAS obtained weights and genotypes of individuals in the two test cohorts:

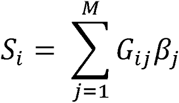

for individual i, the score S_i_ is the product sum of genotypes G_ij_ and weights β_j_ for M SNPs. To make PHS and PRS comparable, we used the identical pruning and clumping process to select independent SNPs for generating the scores. The parameters include clumping within 250kb and linkage disequilibrium greater than 0.1, resulting in 251,040 independent SNPs for generating the scores. No p-value thresholds were imposed to avoid using different numbers of SNPs between the PHS and the PRS. Men-derived scores used weights for SNPs based on the GWAS of men in ADGC, and similarly women-derived scores only used weights from GWAS of women in ADGC. Both men-and women-derived scores were then computed for each participant in the validation cohorts using the same autosomal SNPs. Crossover analyses can thus be used to compare the predictive performance of sex-matched vs. sex-mismatched scores in the validation cohorts.

### Statistical analysis

We implemented genome-wide Cox regression for efficiently estimating hazard ratios across millions of SNPs. P-values of the Cox regressions were obtained using score tests ^34^. The logistic regression GWAS were performed using PLINK. All genome-wide analyses were done using ADGC data, separately for men and women. In order to provide an intuitive interpretation on the obtained weights, we also calculated gene-based effect sizes using Pascal ^35^. Pascal obtained gene-based p-values are based on a linkage-disequilibrium weighted average of effect sizes of SNPs located within 50Kb regions of the gene body.

In NACC, we used 1). Cox regression to examine the predictive power of polygenic scores on AD age-at-onset, and 2). linear mixed effects model to examine the associations between polygenic scores and rate of clinical progression, defined as changes in Cognitive Dementia Rating – Sum of Boxes (CDR-SB). All models were controlled for *APOE* status (dosages of e2 and e4) and education levels. The main analysis of NACC included 2628 men and 3448 women. We also examined whether the patterns of association remained constant if we restricted analyses to neuropathologically-confirmed cases; 817 men and 706 women from NACC had post-mortem neuropathological examinations. To ensure the consistency of the units, all results are based on standardized polygenic scores, comparing changes in 1 standard deviation (SD) of scores.

In ROSMAP, we analyzed the relationship between the neuropathological burden at autopsy and sex-dependent polygenic scores. Four quantifications of neuropathology were included, i.e., the percentage area occupied by β-amyloid, and the density of tau-positive neurofibrillary tangles. Because those neuropathological measures were skewed, we performed a square root transformation to normalize the neuropathology data. We also determined Braak stage, and Consortium to Establish a Registry for Alzheimer’s disease (CERAD) score. All regression models controlled for *APOE* status (dosages of e2 and e4), age-at-death, and education level. To ensure the consistency of the units, all results are based on standardized polygenic scores and neuropathological data, comparing neuropathological variations in 1 standard deviation (SD) of scores.

### Code availability

The code for the genome-wide Cox regressions will be available on GitHub

### Data availability

The summary statistics for genome-wide hazard estimates and gene-based analyses will be found in the Supplemental Materials.

### Distribution of hazard weights

Firt, we performed genome-wide Cox regressions for AD age-at-onset on ADGC individuals (men/women = 7158/10697). The models were controlled for first 5 genetic principal components, *APOE* status, and recruiting sites (Methods). We noticed that there are different top hits between men and women (Figure 2A. and Fure 2B.). Men had a GWAS-significant locus on 1q32.2, encompassing CR1, and women had a GWAS-significant locus on 2q14.3, encompassing BIN1. In addition to GWAS-significant loci, polygenic signals below the GWAS-significant threshold are important for deriving polygenic scores. To provide an intuitive summary on the sex-dependent polygenic effects, we performed gene-based analyses using Pascal ^35^. Figure 2C illustrates the sex-dependent distributions from gene-based analyses. Gene clusters on 19q13.32 continue to show consistent effects between men and women, with trends for sex-specific genetic effects. For example, the effect sizes of *BIN1, MS4A6A, DNAJA2*, and *FERMT2* are larger among women while *FAM193B, C2orf47, TYW5* have larger effect sizes among men. Additionally, the tau-related gene, *MAPT*, shows stronger effects on men than on women.

**Figure 2.**
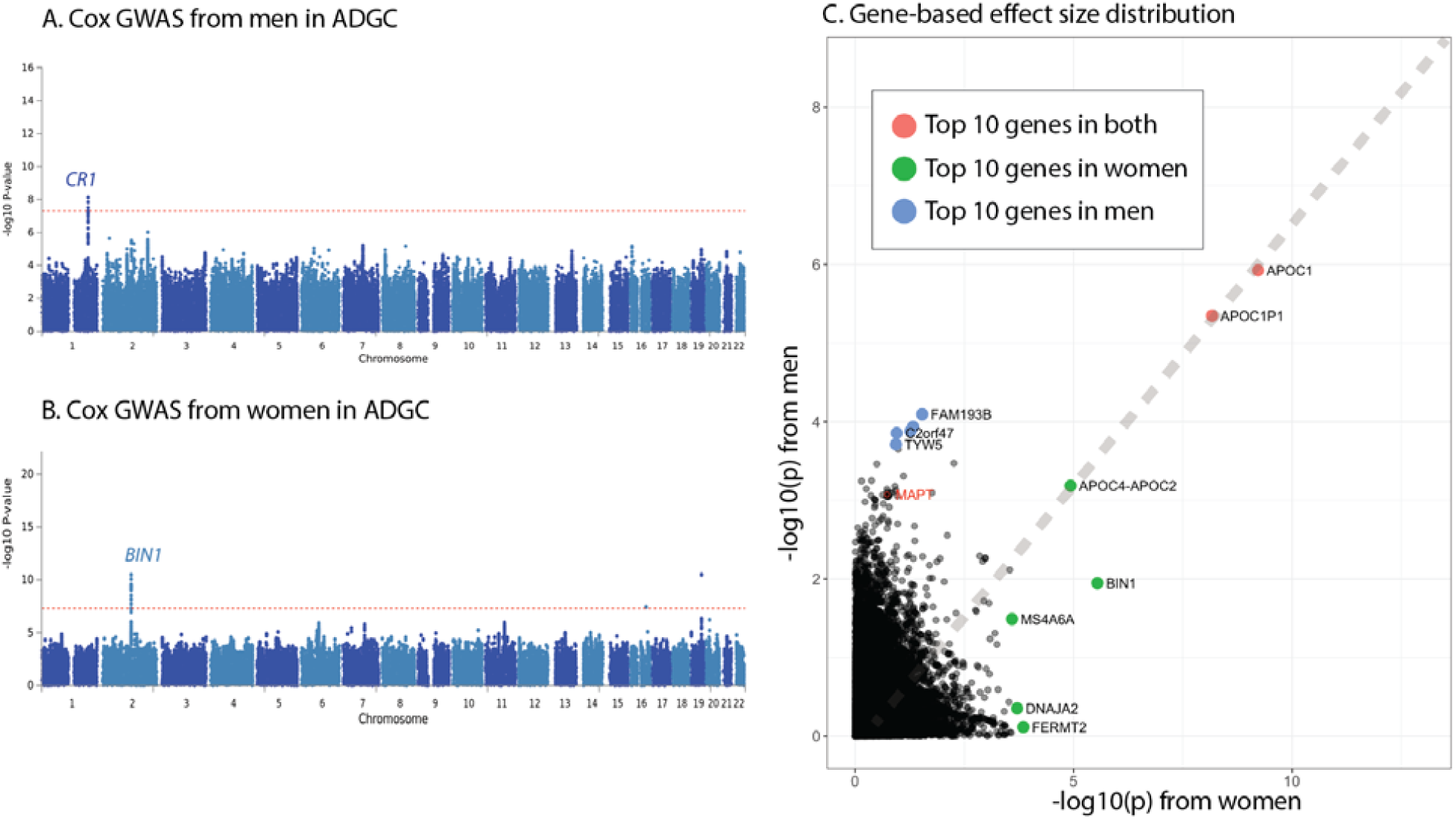
Effect size distributions of obtained hazard weights from sex stratified genomewide Cox regressions. A. Manhattan plot from genomewide Cox regression from men in ADGC. B. Manhattan plot from genomewide Cox regression from women in ADGC. C. Results from gene-based analysis. The diagonal dashed line represents the equivalent effect sizes given the sample size differences. We listed top 10 rank genes in terms of −log10(p) from the Pascal. Genes in both top 10 rank list of men and women are colored in red. Genes in only top 10 rank list of women are colored in green and of men are colored in blue.

### Predicting clinical manifestations in NACC

By aggregating the hazard weights obtained from genome-wide Cox regressions of ADGC, we derived women specific polygenic hazard scores (wPHS) and men specific polygenic hazard scores (mPHS), using standard pruning and clumping process (Methods), for every individual in the NACC cohort (men/women = 2628/3448), resulting in sex-matched model (men with mPHS and women with wPHS) and sex-mismatched model (men with wPHS and women with mPHS). To avoid the confounding of *APOE* due to imputations, we excluded any genetic variants located at *APOE* region (Methods). For clinically determined AD onset, the sex-matched model consistently performed more accurately than the sex-mismatched model (Figure 3A). After controlling for *APOE* status, sex-matched PHS has a hazard ratio (HR) of 1.26 (95% CI: 1.26 – 1.32, p < 1e-16) and sex-mismatched PHS has a hazard ratio (HR) of 1.14 (95% CI: 1.09 – 1.19, p = 1e-10). Sex-matched PHS performed significantly better than sex-mismatched PHS (p = 0.001). Subgroup analyses indicate that stronger predictive power in sex-matched models than sex-mismatched models is evident for both men and women (Supplemental Figure 1A). When we limited our analysis to those with neuropathological disease confirmation (n = 1523), the crossover effects were consistent (HR: 1.21, p = 2e-9, Figure 3B, Supplemental Figure 1B) and retaining significant difference between sex-matched and mismatched models (p = 0.008). Figure 3C shows the performance of polygenic scores in predicting clinical progressions as CDR-SB changes during longitudinal follow-up in NACC. Sex-matched PHS was predictive of annual changes of CDR-SB (β: 0.057, 95% CI: 0.049 – 0.064, p < 1e-16) and performed better than sex-mismatched PHS (β: 0.043, 95% CI: 0.035 – 0.050). The difference between sex-matched PHS and sex-mismatched PHS was statistically significant (p = 0.006). In contrast, PRS from logistic regressions showed no significant associations regardless of which sex-dependent PRS were applied (Figure 3A-C).

**Figure 3.**
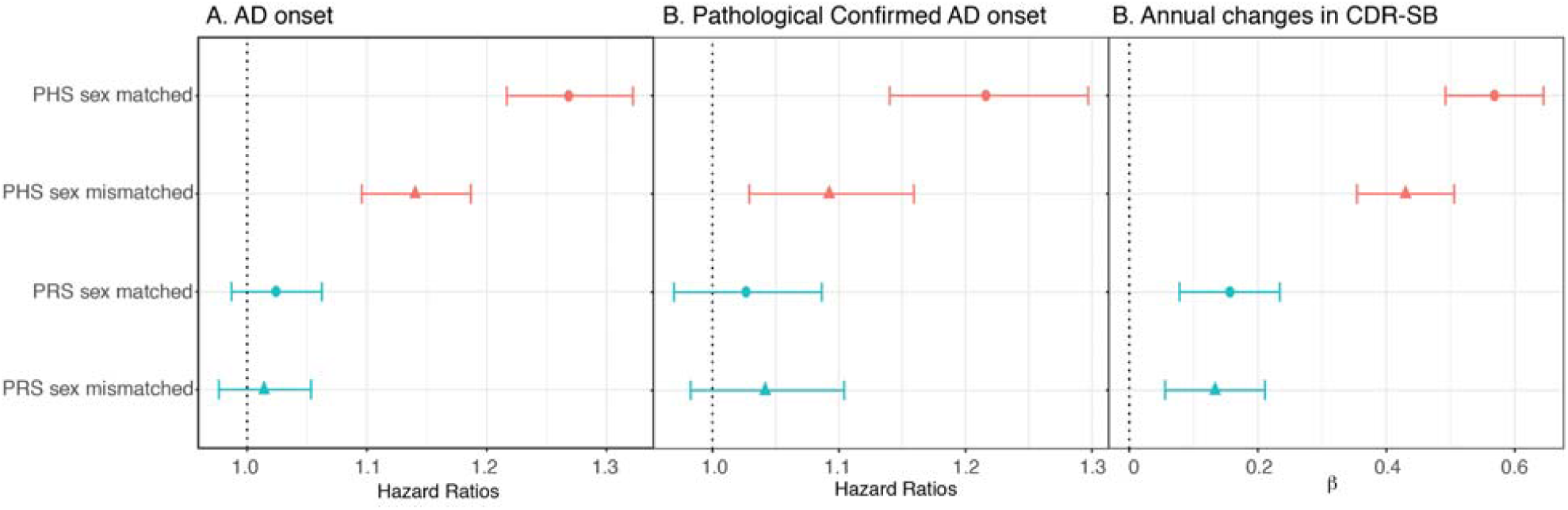
Predictive performance of polygenic components in NACC. Weights from Cox regressions of training data were applied to all participants in NACC, yielding both mPHS and wPHS for all. The hazard ratios of comparing 1 standard deviation differences in PHS, after controlling *APOE* and education levels, are shown. A. Prediction of clinically defined AD. B. Prediction in neuropathologically confirmed AD cases, C. Prediction in CDR-SB changes

### Predicting neuropathology in ROSMAP

Figure 4 demonstrates the association strengths across four different types of neuropathology. After controlling for age at death, education levels, and *APOE* status, sex-matched models have significantly stronger associations than sex-mismatched models for all neuropathological measures (p values for differences between sex-matched and sex-mismatched PHS as 5e-5, 4e-7, 0.007, and 5e-4 for amyloid deposition, CERAD score, tau associated neurofibrillary tangles, and Braak score, respectively). None of the sex-mismatched models reached statistical significance in predicting neuropathology based on polygenic components. Table 2 summarizes the variance explained for subgroup analyses on each neuropathology. Compared to sex-mismatched models, wPHS applied to women increased the variance explained 6 percent, 5 percent, 3 percent, and 6 percent for amyloid deposition, CERAD score, neurofibrillary tangles, and Braak score, respectively. Applying mPHS to men would increase 1 percent, 3 percent, 3 percent, and 4 percent for amyloid deposition, CERAD score, neurofibrillary tangles, and Braak score, respectively. In general, variance explained attributable to the polygenic components for sex matched models can reach up to 89 percent of variance explained by *APOE* only. In contrast, sex-matched PRS had no significant associations with any neuropathologies except CERAD score, whereas no evident sex differences after controlling for *APOE* (Figure 4 and Supplemental Figure 2).

**Table 2.** Variance explained of neuropathological indices for subgroup analysis in ROSMAP.

| Pathology |  | Testing Subjects | Covariates only | plus E2 + E4 | Sex-matched PHS | PHS / APOE * |
| --- | --- | --- | --- | --- | --- | --- |
| **Amyloid Related Pathology** | Amyloid | Women | 2% | 12% | 17% | 55% |
|  | Amyloid | Men | 5% | 12% | 13% | 11% |
|  | CERAD | Women | 1% | 11% | 16% | 51% |
|  | CERAD | Men | 3% | 9% | 12% | 59% |
| **Tau related pathology** | Tangles | Women | 2% | 15% | 19% | 24% |
|  | Tangles | Men | 4% | 10% | 13% | 54% |
|  | Braak | Women | 5% | 11% | 17% | 89% |
|  | Braak | Men | 8% | 15% | 19% | 59% |

\* Amount of variance explained attributable to polygenic component over the amount attributable to *APOE* dosages

**Figure 4.**
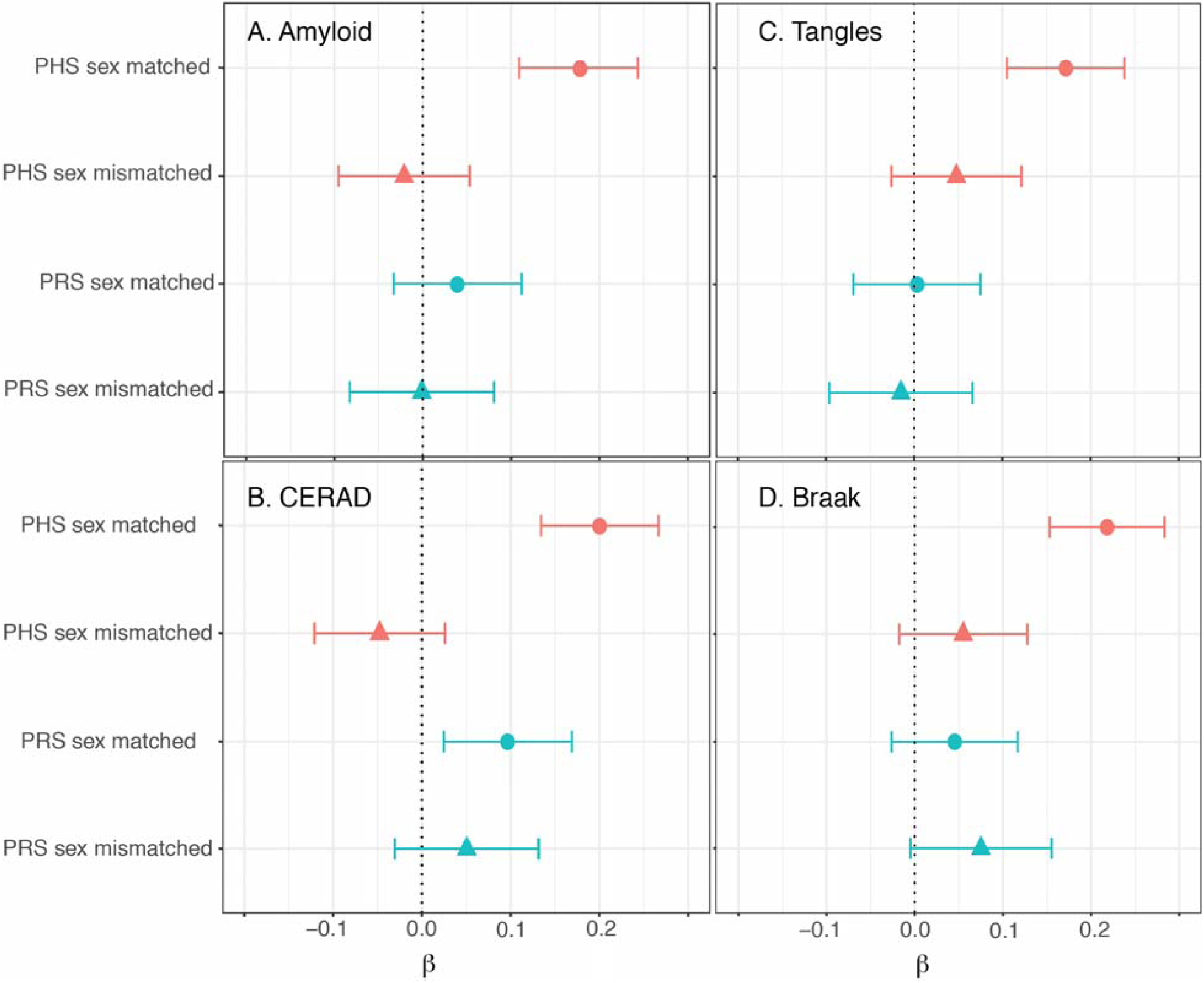
Associations with neuropathology in ROSMAP. Sex dependent polygenic scores were obtained for all participants in ROSMAP. The coloring schemes are consistent with Figure 3. All models controlled for age at death, education levels, and *APOE* status. A. Associations with amyloid deposition, B. Associations with CERAD score, C. Associations with neurofibrillary tangles, D. Associations with Braak score.

### Understanding the sex-dependent polygenic architecture of AD

By modeling the disease courses as time-to-clinical-onset, the polygenic hazard approach revealed sex-dependent autosomal effects on AD after controlling for *APOE*. Sex-matched PHS showed better prediction of both clinical age-at-onset and neuropathological manifestations than sex-mismatched PHS, implying that genetic risk factors differ between men and women. These finding have implications not only for the etiology of AD, but also a new approach to examine sex differences in genetic risks.

Many of the genes highlighted by our analyses have been implicated in AD in prior reports ^28–30,36^. Yet, our survival analyses revealed a complex landscape of sex-dependency across the genome. Loci such as *BIN1, MS4A6A, DNAJA2*, and *FERMT2* contribute higher risk to women than to men. Previous GWAS have identified *BIN1* and *MS4A6A* as risk loci for AD ^36^, but our results indicate that their effects may be sex dependent, especially for pathologica aging processes. Experimental studies have found that *FERMT2* is associated with amyloid deposition^37^ whereas *DNAJA2* interacts with protein tau aggregation ^38^. When aggregating those differences as PHS, the sex-dependency of the genetic effects emerged, indicating there are divergent pathological pathways between men and women.

In addition to the pathogenesis of AD, these crossover analyses also highlight an important aspect for modeling genetic risks – time. AD is an insidious, progressive disease. When the genetic effects on disease risks are differentially expressed across time, the mean liability model cannot readily capture differences in the underlying genetic risks ^39,40^. In our analyses, PRS had limited predictive accuracy on both AD onset and neuropathology, regardless of sex-dependencies. This strongly suggests that explicit modeling of time of clinical disease onset using survival analyses is needed to reveal sex-dependent effects in polygenic signals. Considering one of the key differences between men and women with respect to AD is the temporal disease course, and hence the underlying hazard function, sex-dependent polygenic effects may largely modulate the temporal disease course for AD.

Sex differences are ubiquitous in human biology and disease manifestations, yet are rarely reported in terms of genetic risks ^9^. Our results indicate that by explicitly modeling age-dependent hazards in sex-stratified analyses, we can reveal these sex-dependent effects. In addition to providing insight about sex-differences in AD pathophysiology, we also hope this study will encourage improvements in GWAS study design to consider sex differences regarding time of disease onset.

## Acknowledgement

The work was supported by National Institutes of Health Grants (1R03AG063260-01), National Institute of Aging grants (P30AG10161, R01AG15819, R01AG17917). Acknowledgement for ADGC and NACC can be found in the Supplemental Information.

## Author Contributions

C.C.F., S.J.B., R.D., and A.M.D. conceived and designed the study. C.C.F., S.J.B., and R.D. acquired, analyzed, and interpreted the data. C.C.F., S.J.B., R.D., and A.M.D. drafted the manuscript. W.K.T., C.H.C., L.K.M., C.H.T., W.K., D.A.B., L.A.F., R.M., G.D.S., and O.A.A. critically revised the manuscript for important intellectual content.

## Competing Interests

C.C.F. is under employment of Multimodal Imaging Service, dba Healthlytix, in addition to his research appointment at the University of California, San Diego. A.M.D. is a founder of and holds equity interest in CorTechs Labs and serves on its scientific advisory board. He is also a member of the Scientific Advisory Board of Healthlytix and receives research funding from General Electric Healthcare (GEHC). The terms of these arrangements have been reviewed and approved by the University of California, San Diego in accordance with its conflict of interest policies.

